# Counting particles could give wrong probabilities in Cryo-Electron Microscopy

**DOI:** 10.1101/2025.03.27.644168

**Authors:** Luke Evans, Lars Dingeldein, Roberto Covino, Marc Aurèle Gilles, Erik Thiede, Pilar Cossio

## Abstract

Cryo-electron microscopy (cryo-EM) experiments take 2D snapshots of individual proteins. In principle, these snapshots contain not only the main biomolecular conformations but also scarcely populated states and rare transitions between intermediates. This makes cryo-EM a powerful tool, not only for investigating the structure of biomolecules at high resolution but also for inferring the entire conformational ensemble distribution. Some recent works have reported conformational state populations by counting particle-images from cryo-EM. We wish to caution the community that these measurements are highly susceptible to noise and should not be relied upon as a precise estimate of the thermodynamic landscape of a biomolecule for understanding its biological function. Here, we demonstrate that the extremely noisy nature of cryo-EM images and uncertainty in the viewing orientations of biomolecules lead to ambiguities when assigning images to structures. If ignored, this ambiguity can introduce inherent bias when determining the populations of conformational states through individual particle assignment. We further show that modeling the conformational probability distribution using the entire image dataset mitigates these biases.

## Main

Biomolecules continuously undergo conformational changes within the cell. The population of a given conformation refers to its relative frequency within an ensemble of states. Determining these populations is crucial for understanding how biomolecules achieve their biological function. Several recent studies reported populations from cryo-electron microscopy (cryo-EM) using particle counts assigned to 3D classes or latent-space histograms. We caution the cryo-EM community that these estimates can be distorted by noise and uncertainty, and therefore should not be used as quantitative evidence of the thermodynamics of a biomolecule. However, reliable populations can be recovered by modeling the noise estimated across the entire dataset.

Populations reflect the probability density of conformations, and recovering it from data is an inverse problem: we observe an ensemble of images 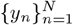 sampled from the data distribution *p*_data_(*y*), and attempt to recover a probability density *π*(*x*) over conformations *x* that generated the images. The population forward model is obtained by convolving the conformational probability density with the imaging forward model, *p*(*y*|*x*), also known as the likelihood, which results in a convoluted probability density for the observed data,

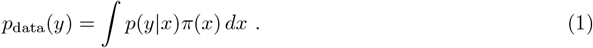

Counting and histogramming approaches estimate *π*(*x*) by tallying up particle assignments and organizing them into specific classes or locations in an underlying space using the likelihood *p*(*y*|*x*) (Supplementary Note). But in the presence of experimental noise, images could potentially be assigned to multiple conformations as the likelihood *p*(*y*|*x*) exhibits high variance due to the high noise levels. This makes it challenging to assign an image to a specific conformation with high precision. Choosing a conformation requires knowledge of *π*(*x*), which requires solving the inverse problem (1) [5]. If one assigns an image without this knowledge, one commits the “base-rate fallacy”: a statistical fallacy of ignoring global statistics when making inferences on a single sample [6].

We illustrate this failure mode of particle counting on a discrete two-state system: the spike protein in two conformations (Figure 1a). First, we prepared several particle datasets with varying noise levels, all with a ground truth population of 80% particles in the 1-up state and 20% in the 3-down state (Supplementary Note and Figure 1b). We used traditional 3D classification techniques (yellow) that attempt to classify which 3D structure each image belongs to. Even with ground truth poses, these techniques fail quickly, giving incorrect population estimates as noise is increased. Hard and soft assignment (blue and green, respectively) show the same trend. In contrast, methods that explicitly attempt to solve the inverse problem (1), *e*.*g*., by ensemble reweighting [2] or by deconvolution [3], give accurate populations even at high noise levels. We observed similar trends in heavily classified real images (*<* 10% of the initial picked particles in EMPIAR-12098), which we prepared in an 80/20 mixture of the 3-up and 2-up spike states, respectively. With available alignments, all methods performed relatively well, though particle-counting approaches showed slightly lower accuracy (Figure 1c, left). However, in a typical workflow, particle poses and contrast scaling are often uncertain and require refinement. We exemplified this uncertainty by adding noise to the images. In this scenario, the accuracy of particle counting methods decreased, while the ensemble reweighting and deconvolution retained the correct population estimates (Figure 1c, right). These results confirm that 3D classification can be inconsistent and unstable [7] due to significant uncertainty in assigning individual images to specific conformations.

**Figure 1.**
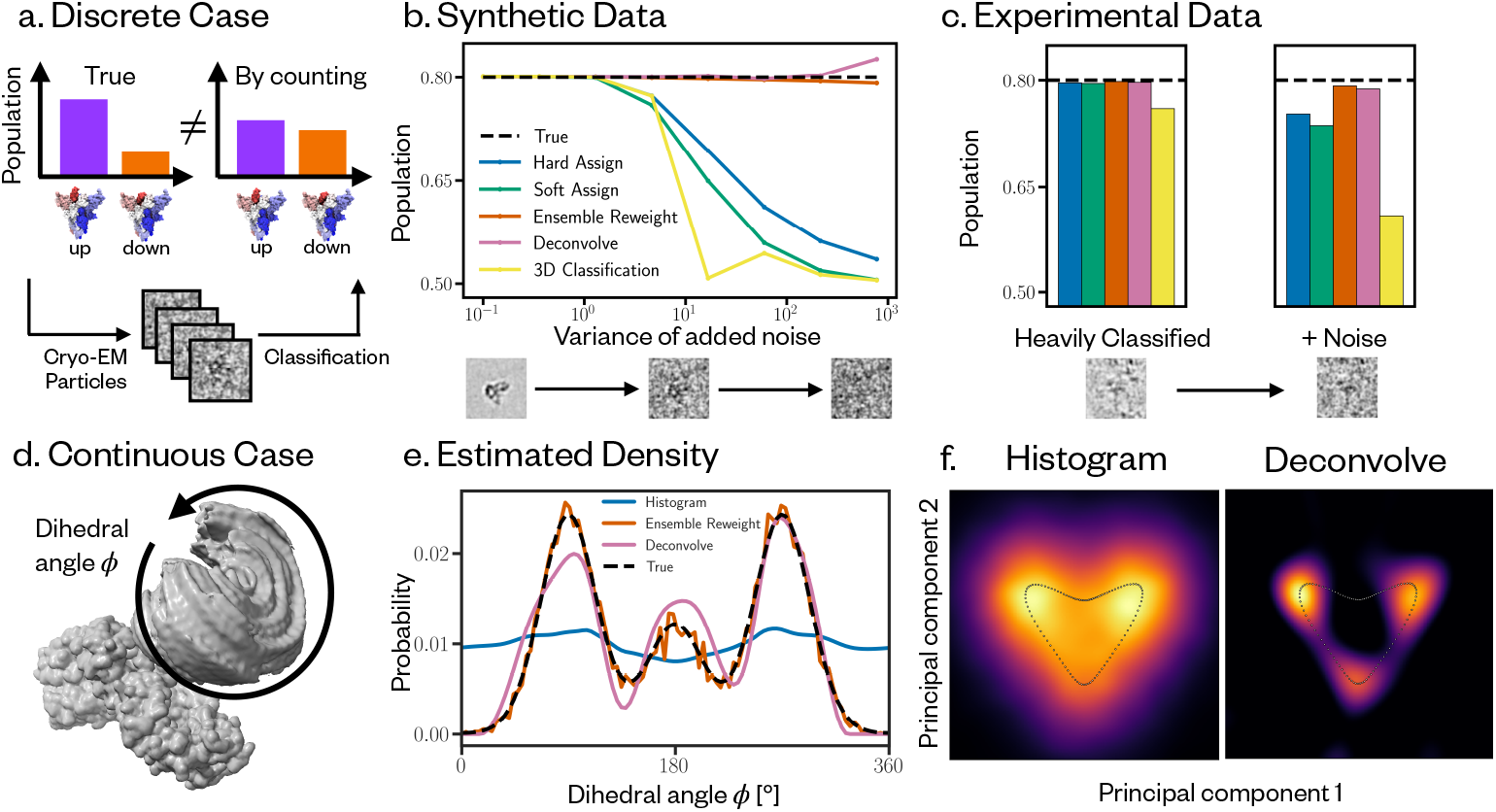
(a) The concept motivating our correspondence: counting cryo-EM particles can fail to recover the population of classes under high noise. Estimated population of the spike protein in the 1-up state for synthetic datasets with increasing added noise (b) and in the 3-up state for real data (EMPIAR-12098) and added noise (c) for hard (blue), soft (green) assignment, 3D classification (yellow) [1], ensemble reweighting [2] (orange) and deconvolution [3] (pink). The true population is shown as a dashed line and plotted below are low-pass filtered images at representative noise levels. IgG-1D domain from cryoBench [4] reconstructed using RECOVAR [3] with particles sampled from a multimodal distribution along the dihedral angle. (e) Ground truth probability density (dotted black line) along the dihedral angle, compared against the estimates from the histogram (blue), the deconvolved (pink), and the ensemble reweighted (orange) densities. (f) Probability density along the first two principal component assignments of images by histogramming (left) and by deconvolution (right). The black dots corresponds to the location of the ground truth volumes. Details about the datasets and methods are provided in the Supplementary Note.

Continuous heterogeneity methods have recently shifted the cryo-EM community from discrete classification to latent-space embedding of individual images. However, latent-space distributions are still highly affected by noise. Inspired by the Flatiron Institute 2023 cryo-EM heterogeneity challenge [8], we showcase this on a simple example: the rotation of the IgG domain from cryoBench [4], where the underlying conformational density is described along one dihedral angle (Figure 1d). Here, we sample a multimodal distribution on the dihedral (Figure 1e, black dashed line) and compute images at high noise (Supplementary Note). We take RECOVAR [3], a high performing method in cryoBench, and assign particles to latent coordinates defined by the first two principal components. Latent-space analysis in continuous heterogeneity methods is typically performed by creating a histogram of the embedded image assignments (Figure 1f-left). This predicts a nearly flat distribution, entirely missing the middle mode (blue - Figure 1e). In contrast, solving the inverse problem (1) can be performed by denoising the latent space using the deconvolution approach [3] (Figure 1f-right). This postprocessing correctly finds all modes of the distribution (pink - Figure 1e). If the conformations are given, the ensemble reweighting method [2] accurately infers the correct probability distribution (orange - Figure 1e).

Here, we demonstrated that noise-induced bias can affect population estimates when counting cryo-EM particles. This bias depends on the level of noise and uncertainty in the particle images as well as the magnitude and distinguishability of the conformational changes. At low noise levels, counting or histogramming can provide reliable population estimates (Figure 1b-left). However, determining the exact noise regime of real data is not possible, and high noise and uncertainty can hinder the accurate recovery of probability distributions through 3D classification or latent space histogramming. Therefore, scientists should exercise caution when drawing biological conclusions based on classification or latent-space assignments in cryo-EM. In contrast, methods that account for noise and uncertainties by leveraging the statistics of the entire particle stack [2, 3] yield more reliable estimates, though they remain susceptible to failure and highlight the need for further research.

## Acknowledgments

The Flatiron Institute is a division of the Simons Foundation. L.D. and R.C. acknowledge the support of Goethe University Frankfurt, the Frankfurt Institute of Advanced Studies, the LOEWE Center for Multiscale Modelling in Life Sciences of the state of Hesse, the CRC 1507: Membrane-associated Protein Assemblies, Machineries, and Supercomplexes (P09), and the International Max Planck Research School on Cellular Biophysics. L.D., and R.C. thank the Flatiron Institute for hospitality while a portion this research was carried out. E.H.T. was supported by Cornell University. We thank Niko Grigorieff, David Silva-Sanchez, Minhuan Li, Robert Gower, Ellen Zhong and Sonya Hanson for feedback and insightful discussions.

## Supplementary Note

## 1 Counting particles in Cryo-Electron Microscopy

### 1.1 Image-to-structure likelihood

We consider a weak phase approximation for cryo-electron microscopy (cryo-EM) image formation [9–11]. We represent a conformation 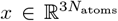 by its electron density. An image *y* is formed from applying a 3D rotation to the electron density, then a projection along the *z* axis, a convolution with the point-spread function and then a translation in the *xy* plane. Let Θ represent the parameters for the translation, rotation and point-spread function, and ℱ (Θ, *x*) represent this forward model of the image formation process. Assuming pixel i.i.d. Gaussian white noise with standard deviation *λ*, the likelihood of an image given a structure is

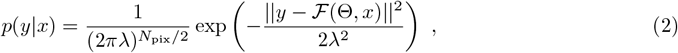

where ||·||^2^ is the *l*^2^-norm and we have assumed all Θ parameters are known, e.g. previously estimated from *ab initio* reconstruction. In essence, the likelihood in Eq. (2) is the one used for 3D classification, being the key element for associating particles with classes in cryo-EM.

### 1.2 Cryo-EM particle assignment

Eq. (2) gives the probability of seeing an image given a specific structure. While classification algorithms and software vary considerably, most approaches utilize the likelihood to assign images to structures [1, 5, 9, 12].

We focus here on the case of discrete classification. However, we note that this can be easily generalized to continuous spaces as shown in section 2.2. In discrete classification, we assume that each image *y*_*i*_ is generated by one of *M* classes, with conformations *x*_1_, …, *x*_*M*_. A class assignment for image *y*_*i*_ is a label *z*_*i*_ ∈ {1, …, *M*} associated to the class 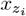

#### 1.2.1 Hard particle assignment

An intuitive, but subtly incorrect, classification approach is to assign each image *y*_*i*_ to the class with the highest likelihood. Denoting the label to which we assign image *y*_*i*_ as *z*_*i*_, the *likelihood-based hard assignment* is defined as

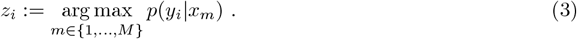

One can then attempt to estimate the population of each class by counting the number of images assigned to each class. Therefore, the estimated population for class *m*, denoted *p*_hard_(*x*_*m*_), is defined as

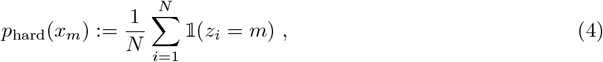

Where 1 (*z*_*i*_ = *m*) = 1 if *z*_*i*_ = *m*, and 0 otherwise. Eq. (4) is the population estimate used for the “hard assign” examples in the Main Text (blue, Figure 1b-1c Main Text), which illustrate that likelihood-based hard assignment will not recover well the correct populations at low SNR. Note that we can interpret hard assignment exactly as the *assignment step* for k-means clustering [13] on centroids {*x*_*m*_}.

#### 1.2.2 Soft particle assignment

While simple, hard-assignment ignores any uncertainty in the assignment of images to classes: it is quite possible that multiple conformations have a reasonable chance of generating a given image. Soft assignment addresses this problem by assigning a probability to each class for each image. In *likelihood-based soft assignment*, images are assigned proportionally to the likelihood of each class, *i*.*e*.,

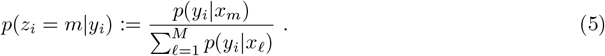

The population of each class is the average of these posterior probabilities over all images [5, 9, 12, 14]:

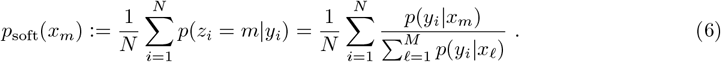

Eq. (6) is the population estimate used for the “soft assign” examples in the Main Text (green, Figure 1b-1c Main Text), which illustrate that likelihood-based soft assignment will not recover well the correct populations at low SNR.

Analogously with hard classification and k-means, the class probabilities in Eq. (5) are *responsibilities* in expectation maximization terminology, and the class population defined by Eq. (6) is the *M-step* for mixture weights in expectation maximization [13].

### 1.3 Particle assignment using the joint probability

While intuitive, the likelihood-based hard and soft assignments introduce a systemic bias into the estimated class probabilities. In Bayesian classification frameworks used specifically in cryo-EM [5, 9] and generally in machine learning [13, 15, 16], one uses the *joint probability* rather than just the likelihood to classify data.

This approach assumes that a class *x*_*m*_ has a probability *π*(*x*_*m*_) := *π*_*m*_ to occur, so that a list of classes *x*_1_, …, *x*_*M*_ has probabilities *π*_1_, … *π*_*M*_ with 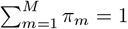. The joint probability density gives the probability of observing the image *y*_*i*_ while a molecule adopts class *x*_*m*_, and is given by

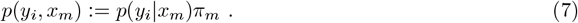

Given *π*, the posterior probabilities from Eq. (5) are generalized to

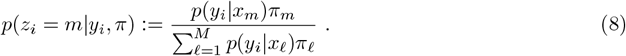

Depending on the inference or classification problem, *π* may be user-chosen [13, 15, 16], but in other cases, one can *infer* the *π* that fits best with the image dataset {*y*_*i*_} [5, 17].

#### 1.3.1 Comparing assignments using the joint probability or likelihood

Comparing with Eq. (5), we see that the posterior from likelihood-based assignment assumes that *π* is uniform with *π*_*m*_ = 1*/M*. This introduces a bias into the resulting assignments because the baseline (true) probability *π*^*^ has not been taken into account. The synthetic spike example and IgG example (Figures 1a-b, 1d-f Main Text) are direct examples of this: if the true conformations {*x*_*m*_} follow a probability distribution *π*^*^ and the images have low SNR, counting from a classification with uniform *π* will be incorrect, often drastically so, even when alignments and CTF are estimated perfectly.

### 1.4 Differences in Cryo-EM classification approaches

A comparison of cryo-EM classification methods is beyond the scope of this work. However, we wish to emphasize that existing literature varies significantly between approaches, and the gap between published papers and their corresponding software can often be unclear. Some classification approaches use expectation maximization, but differ in what parameters they update (Eq. 25 of [5], Eq. 6 of [18]), while others define *π*_*m*_ = 1*/M* for *m* = 1, …, *M* and use alternative optimization schemes (supplementary Eq. 6 in [1]). Related work has indicated that 3D classification can inconsistently estimate poses and classifications [7]. We highlight that studies on the robustness of methods for heterogeneity analysis [4, 8] can greatly benefit the cryo-EM community.

## 2 Explicitly estimating the conformational probability density

To solve the inverse problem (Eq. 1 Main Text) of matching a proposed conformational distribution *π*(*x*) to an observed single particle image dataset, a natural approach is to see which proposed *π*(*x*) is most statistically likely to have generated the observed data according to a forward imaging model *p*(*y*|*x*), such as the image-to-structure likelihood in Eq. (2).

### 2.1 Ensemble Reweighting

The ensemble reweighting method [2] employs Bayesian analysis to infer *π*(*x*) from individual cryo-EM images. Suppose we have a candidate distribution *π*(*x*) over biomolecular conformations, which can be defined by a set of parameters that we intend to infer. Then, using the imaging forward model *p*(*y*|*x*), the marginal likelihood of observing an image *y* given distribution *π*(*x*) is the average over all possible conformations *x* which could have been generated from distribution *π*,

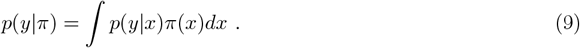

Given single particle images *Y* = {*y*_1_, …, *y*_*N*_} sampled i.i.d, the marginal likelihood of observing the dataset *Y* given the distribution *π* is

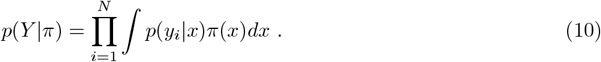

To incorporate uncertainty in the distribution *π*, one can define a prior *p*(*π*) and take the Bayesian approach of sampling the posterior *π* ∼*P* (*π*|*Y*) ∝ *p*(*Y*|*π*)*p*(*π*), further described in ref. [2]. In practice, how one actually evaluates *P* (*Y*|*π*) from Eq. (10) for a given *π*(*x*) heavily depends on how it is parametrized. The discrete case of ensemble reweighting [2] and classification [5, 19] corresponds to *π*(*x*) = _*m*_ *π*_*m*_*δ*(*x*− *x*_*m*_) for a fixed set of structures {*x*_*m*_} with weights {*π*_*m*_}, resulting in the marginal likelihood

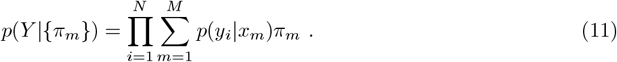

Because Eq. (11) directly uses the values of the probability distribution *π* at the structures {*x*_*m*_}, inferring *π* from Eq. (11) is an example of a *non-parametric maximum likelihood estimation* problem [17, 20, 21].

The main objective of the ensemble reweighting method is to estimate {*π*_*m*_} from the entire set of images, instead of from individual assignments, by using the likelihood from Eq. (2). The method takes the image-to-structure likelihood *p*(*y*_*i*_|*x*_*m*_) at all images and structures (the same input as for hard or soft classification) and outputs the weights {*π*_*m*_} calculated by directly maximizing Eq. (11) or by a Bayesian approach with Markov chain Monte Carlo (as in ref. [2]). For ensemble reweighting in this work, we optimize the weights {*π*_*m*_} to maximize the marginal likelihood in Eq. (11). We use an iterative scheme, where the weights at step *t*, denoted 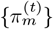, are updated as

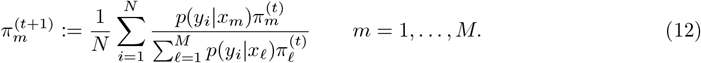

and repeated until convergence. Eq. (12) is the population estimate used for the “ensemble reweight” examples in the Main Text (orange, Figure 1Main Text). In practice, we stop iterating when

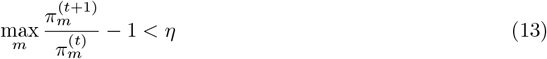

for a chosen threshold *η*. This convergence criteria follows directly from the convexity of the log marginal likelihood [22, 23], particularly that

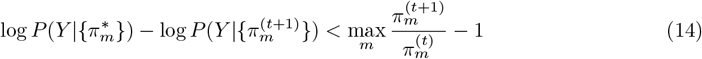

for an (unknown) optimal solution 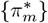, and hence the threshold *η* guarantees a marginal likelihood within *η* of the true maximum marginal likelihood.

Indeed, the update in Eq. (12) is a generalization of the population estimate from soft assignment in Eq. (5): we soft assign with class prior 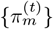 and then average to obtain class populations 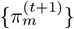. This is precisely the class probability update in multi-class expectation maximization as performed in FREALIGN with fixed volumes and alignments [5], and is analogous to the mixture weight update for expectation maximization on mixture models [13]. Historically, the exact iteration scheme in Eq. (12) has been used in various scientific contexts for over 50 years in astrophysics, finance, information theory, and statistics [17, 22–26]. Our distinction from refs. [5] is that we assume structures have already been estimated, and so we do not update volumes alongside the weights *π*, which poses a more challenging optimization problem with a typically non-convex optimization landscape. This parallel directly illustrates that the ensemble reweighting framework *decouples* the estimation of structures from the inference of the conformational probability density on the structures.

### 2.2 Recovery from deconvolution of latent prediction

An alternate approach to recovering state probabilities, compared to the methods in Section 2.1, would be to explicitly undo the bias introduced by the classification methods. If one directly uses Eq. 1 of the Main Text, the basic idea of deconvolution is to directly estimate the probability *p*_data_(*y*) at each image *y*_*i*_ and then solve for *π* to minimize the error between *p*_data_(*y*) and *p*(*y*|*x*)*π*(*x*)*dx* over {*y*_*i*_}. However, starting from this equation is generally not feasible, as the images are very high dimensional and using a density estimator biases our statistical model. Some of the original literature on expectation maximization compares the method with deconvolution [17, 24], but only for 1-dimensional data.

For the cryo-EM case, one can make the deconvolution strategy tractable by using the latent data embeddings instead of images. This is the basis of the deconvolution strategy presented in ref. [3]. In the discrete case (as in section 1.2-1.3), a latent assignment *z*_*i*_ is a label from classification, with *z*_*i*_ ∈ {1, …, *M*} for *M* classes. In the continuous case, the label is a vector *z*_*i*_ ∈ ℝ_*d*_, where *d* ≪ 3*N*_atoms_, and *z*_*i*_ = *f* (*y*_*i*_) for a feature mapping *f* (·) (*e*.*g*., a neural network or PCA embedding).

First, we assign images to structures according to an assignment *p*(*z*| *y*). For a deterministic assignment, this a delta function *p*(*z*|*y*) = *δ*(*z*−*f* (*y*)), but this can also be probabilistic as in the soft assignment case (Eq. (5)) or for variational inference approaches. This assignment generates a new probability distribution over latent space

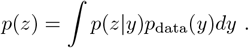

Based on our assumption from Eq. 1 Main Text, we can substitute for *p*_*data*_(*y*) and then change order of integration,

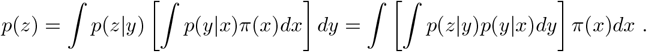

Substituting *p*(*z*|*x*) := *p*(*z*|*y*)*p*(*y*|*x*)*dy*, we have the latent density

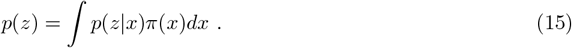

If we can compute *p*(*z*|*x*), we can deconvolve Eq. (15) in *d* dimensions, a significantly reduced dimensionality compared to the image space.

This “deconvolution” algorithm can be summarized as:

1. Use a standard density estimation technique to estimate *p*(*z*) (*e*.*g*., hard-assignment in the discrete case, or a kernel density estimator in the continuous case).
2. Estimate the kernel *p*(*z*|*x*) for a discretization of *x* and *z*.
3. Solve Eq. (15) for *π*(*x*).

Unfortunately, when the embedding is done by a complicated nonlinear map like a neural network, computing the kernel *p*(*z*|*x*) directly is typically prohibitive to compute. Still, there are important scenarios in which *p*(*z*|*x*) can be computed easily, making the estimation of *π*(*x*) computationally tractable. We illustrate this fact with an example for the discrete case here.

Specifically, we consider the case of estimating the relative density of two states *x*_1_ and *x*_2_ from a dataset with known poses and known classes (*i*.*e*., *x*_1_ and *x*_2_ are known in advance) as in Fig. 1a Main Text. In this case, *z* can only take the values 1 or 2, and the integral in Eq. (15) evaluated at *z* = 1 reduces to a sum of two terms

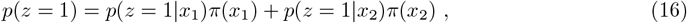

where *p*(*z* = 1|*x*_2_) is the *missclassification rate* (*i*.*e*, the proportion of images of *x*_2_ which are misclassified as *x*_1_). In other words, Eq. (16) states that the proportion of images predicted to be of class *x*_1_ is the proportion of class *x*_1_ times the correct classification rate, plus the proportion of images from class *x*_2_ times the missclassification rate.

For a given parameter Θ_*i*_ (poses, CTF, etc.), the misclassification rate can be readily computed analytically by finding conditions where

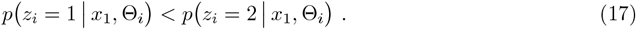

Some algebra shows that this happens when 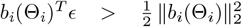 where 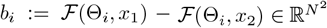. Since *ϵ* is assumed to be white Gaussian noise, 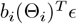 is a standard Gaussian with mean 0 and variance 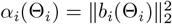. Thus, the missclassification rate corresponds to the integral of the of a standard normal Gaussian, which is: 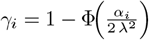 where Φ is the CDF of the standard normal distribution. By integrating over the parameter Θ_*i*_, we obtain the average missclassification over the dataset, *γ* = *γ*(Θ_*i*_)*d*Θ_*i*_.

If *x*_1_ occurs with probability *π*(*x*_1_) and *x*_2_ with probability *π*(*x*_2_) = 1−*π*(*x*_1_), then the fraction of labels predicted to state 1 is

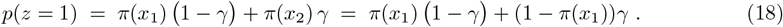

Since *p*(*z* = 1) and *γ* can be estimated from data, the fraction of images coming from image 1 is thus: *π*(*x*_1_) = (*p*(*z* = 1) − *γ*)*/*(1 − 2*γ*).

The procedure can be summarized as follows: first, count the proportion of images hard-assigned to state 1, denoted *p*(*z* = 1); then estimate the misclassification rate *γ* using the formula: 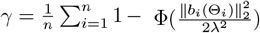, and afterwards estimate the population of state 1 as 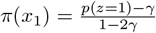.

In the continuous case using a PCA embedding developed in ref. [3], the exact same procedure applies, with only minor modifications needed to handle the continuous nature of the problem: first, a continuous kernel density estimate is formed to estimate *p*(*z*), then a “continuous missclassification” kernel is estimated *γ*(*z, x*) and afterward a numerical scheme is used to solve the resulting system of equations.

The two state procedure above is the population estimate used for the “deconvolve” example in the synthetic and experimental spike examples of the Main Text (pink, Figure 1b-c, Main Text), and the method from [3] is used for the “deconvolve” example in the IgG example of the Main text (pink, Figures 1e-f, Main Text).

## 3 Discrete classification: spike examples

### 3.1 Synthetic data

We first present a synthetic example of how the population estimate of the spike protein with the receptor binding domain (RBD) down versus 1-up can fail in 3D classification for images with large amounts of noise, even when class volumes are optimal. We simulate 10 datasets of different signal-to-noise ratios (SNRs), each dataset with 100,000 images where 80 % of images from 1-up and 20 % from down states, respectively. We note that the structures are identical apart from the position of the RBD. Each dataset was simulated using the forward model in RECOVAR [3] with defocus randomly sampled from EMPIAR-10076 [27], centered images, and uniformly distributed poses in SO3. The pixel size was 1.3°A and the box size was 256 pixels per dimension.

For each given image *y*_*i*_ we compute the likelihood that the image was generated from the 1-up state, *p*(*y*_*i*_ |*x*_up_), and and the likelihood that *y*_*i*_ was generated from the down state *p*(*y*_*i*_|*x*_down_). To emphasize how strongly noise affects particle counting even when no other data processing is misspecified, we compute these likelihoods using the ground-truth imaging parameters with Eq. (2). Using the likelihood values, we compute the populations for hard and soft classification of the images as described above. We compare this with the estimates from ensemble reweighting and deconvolution methods. We also compare these results to those from 3D classification in cryoSPARC [1], inputting the ground truth volumes for Initialization mode: input. For 3D classification, we use a filter resolution of 5°A and all default settings for the cryoSPARC 4.6 version of the job.

For the datasets with low noise level, all methods accurately predict 80% population in the 1-up state (far left of Figure 1b - Main Text). For the datasets with intermediate to high noise, the methods that perform per-particle counting (hard, soft and 3D classification) fail to accurately estimate the population.

### 3.2 Experimental data

We use experimental cryo-EM images from EMPIAR-12098 [28] that were heavily classified (*<* 10% of the initial particles) to reconstruct high-resolution volumes of the spike in the 3-up and 2-up states. Here, we treat the classes as ground truth and create a mixed dataset by combining the 3-up and 2- up depositions. We use the cryoSPARC restack utility to combine all 28976 particles of EMD-50421 (3-up) with 7244 particles from the EMD-50422 (2-up), resulting in 80% particles from the 3-up state and 20% particles from the 2-up. We use RECOVAR to compute reconstructions *x*_3-up_ and *x*_2-up_ from the full EMD-50421 and EMD-50422 particles respectively, and then low-pass filter to 20°A, to mimic the initial round of cryoSPARC 3D classification. We utilize the pose and defocus information from the original EMD-50421 and EMD-50422 data. We Fourier downsample the original images to box size 200 pixels per dimension and pixel size 1.78°A.

To illustrate the sensitivity of particle counting, we create a perturbed dataset by normalizing the mixed dataset to have average image norm 1 and then adding i.i.d Gaussian white noise with variance 10 (see examples in Main Text Figure 1c). We then compute the likelihoods on the two datasets, (*i*) the mixed dataset of the original 3-up/2-up particles and (*ii*) the same dataset with noise added. To compute the likelihoods, we first estimate the noise variance for each dataset using the RECOVAR pipeline. Then, for each image *y*_*i*_ we use the reference volume for that state to calculate the likelihood using the original poses and defocus. With the likelihood values, we compute the populations from hard and soft assignments, and ensemble reweighting and deconvolution as described above. We compare these results to cryoSPARC 3D classification, inputting the *x*_3-up_ and *x*_2-up_ volumes for Initialization mode: input. For 3D classification, we use a filter resolution of 20°A and all default settings for the cryoSPARC 4.6 version of the job. In Figure 1c - Main text, we plot the estimated populations of the 3-up state for both datasets. We show that methods that take into account the full dataset perform better than those that perform individual particle assignments for data perturbed by noise.

## 4 Continuous Heterogeneity: IgG-1D Example

We consider synthetic data from the human immunoglobulin G (IgG) antibody, generated from atomic models used in CryoBench [4]. These models are generated from a ground truth structure (PDB: 1HZH) by rotating a fragment antibody (Fab) domain at 100 different dihedral angles. Using these conformations, we simulate one dataset at low signal-to-noise with 100, 000 images. The images follow a multimodal probability distribution, the von Mises distribution mixture, given in degrees by

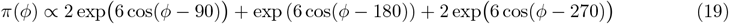

on the dihedral angle *ϕ* (black dashed line, Figure 1e- Main Text). The von Mises distribution [29] is a common approximation to the normal distribution on the circle, and so the distribution in Eq. (19) is analogous to a mixture of Gaussians on the dihedral angle *ϕ* ∈ [0^°^, 360^°^] with peaks at 90^°^ and 270^°^ twice as high than at 180^°^.

We simulated the particles using the forward model in RECOVAR [3] with defocus randomly sampled from EMPIAR-10076 [27], centered images, and uniformly distributed poses in SO3. The pixel size was 1.3°A and the box size was 256 pixels per dimension.

We apply a mask on the Fab domain and then run the RECOVAR [3] pipeline with 20 dimensional embeddings and latent space estimation in 2 dimensions. Here, the histogrammed density (left, Figure 1f - Main Text) is a smoothed histogram by kernel density estimation, with bandwidth selected by Silverman’s rule [30], and the deconvolved density (right, Figure 1f - Main Text) is from the deconvolution step of the RECOVAR pipeline, with the default regularization chosen as described in [3]. The heart-shaped black dots in Figure 1f are the PC coordinates 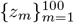 of the ground truth volumes 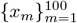.

We take the histogrammed and deconvolved densities evaluated just on the embedded volumes {*z*_*m*_}, namely the density values at black dots of Figure 1f - Main text, and plot the probability density along the dihedral angle in Figure 1e - Main text. The histogrammed density detects only two slight modes and is essentially a flat landscape. In contrast, the deconvolved density detects all three modes and low support regions of the ground truth conformational density. For a reference comparison, we compute the image-to-volume likelihoods from the ground truth volumes and apply the ensemble reweighting method as described in section 2.1. The density has close agreement with the ground truth density, and indicates consistency between the decovonolution and ensemble reweighting methodologies (Figure 1e - Main text).

